# Src42A required for collective border cell migration *in vivo*

**DOI:** 10.1101/186049

**Authors:** Yasmin Sallak, Alba Yurani Torres, Hongyan Yin, Denise Montell

**Affiliations:** Molecular, Cellular, and Developmental Biology Department University of California, Santa Barbara, CA 93106; Department of Biological Chemistry and Center for Cell Dynamics, Johns Hopkins School of Medicine, 725 North Wolfe Street, Baltimore, MD 21205

## Abstract

The tyrosine kinase Src is over-expressed in numerous human cancers and is associated with poor prognosis. While Src has been extensively studied, its contributions to collective cell migration *in vivo* remain incompletely understood. Here we show that Src42A, but not Src64, is required for the specification and migration of the border cells in the Drosophila ovary, a well-developed and genetically tractable *in vivo* cell migration model. We found active Src42A enriched at border cell/nurse cell interfaces, where E-cadherin is less abundant, and depleted from border cell/border cell and border cell/polar cell junctions where E-cadherin is more stable, whereas total Src42A protein co-localizes with E-cadherin. Over-expression of wild type Src42A mislocalized Src activity and prevented border cell migration. Constitutively active or kinase dead forms of Src42A also impeded border cells. These findings establish border cells as a model for investigating the mechanisms of action of Src in cooperative, collective, cell-on-cell migration *in vivo*.

## Introduction

*c-src* is a proto-oncogene that encodes a nonreceptor tyrosine kinase that is overexpressed and/or hyper-activated in many human cancers. Src has been extensively studied, primarily in fibroblasts cultured on coverslips, where it affects cell proliferation, adhesion, morphology, survival, and migration [reviewed in (Yeatman, 2004)]. However, much remains unknown regarding the function of Src in epithelial cells, in complex 3-dimensional environments *in vivo*. So how Src expression and activity contribute to tumor progression and metastasis remains unclear. Recent studies suggest that collective cell migration contributes significantly to metastasis [reviewed in (Cheung and Ewald, 2016; Lambert et al., 2017)]. Src activity is essential for promoting collective invasion of cancer cells (Canel et al., 2010) but Src activity is best known for inhibiting cell-cell and cell-matrix adhesion. So, if Src kinases contribute to collective cell migration, the question arises as to how Src activity is regulated to permit selective maintenance of some cell-cell adhesions and loss of others.

Border cells in the Drosophila ovary serve as a useful and well-studied model of collective cell migration *in vivo* [reviewed in (Montell et al., 2012)]. Each fly egg chamber is composed of 16 germline cells (15 support cells called nurse cells and one oocyte) surrounded by a monolayer of epithelial follicle cells. At the anterior and posterior poles of each egg chamber, two special cells named polar cells develop. Border cell clusters form at the anterior end of the follicular epithelium. Polar cells secrete a cytokine known as Unpaired (Upd), which activates Janus kinase (Jak) signaling in nearby cells, in turn activating Signal Transducer and Activator of Transcription (STAT), which is necessary for border cell specification. Once specified by Jak/STAT signaling, border cells migrate collectively as a group of 4-7 migratory cells surrounding the two non-migratory polar cells. Border cells migrate surrounded on all sides by nurse cells. Therefore, border cell migration serves as a model of cooperative, collective, cell-on-cell migration. In addition to Jak/STAT signaling, border cell migration relies on a complex network of receptor tyrosine kinase (RTK), steroid hormone, and Jun N-terminal kinase signaling, as well as proteins that regulate cytoskeletal dynamics including Rac, Rho, and Cdc42 GTPases. The regulation of E-cadherin-mediated adhesion is particularly critical for collective direction sensing (Cai et al., 2014). Each of these molecular pathways can interact with Src kinases. Therefore, we set out to determine which, if any, Src kinases are required for collective, cooperative, cell-on-cell migration using border cells as a model.

In mammals, *in vivo* loss-of-function studies of Src kinases are hampered by the potential redundancy amongst the 11 Src family kinase members. In contrast, the Drosophila genome encodes only two: Src64 and Src42A. Here we report the requirement for Src42A, but not Src64, in collective migration of the border cells *in vivo*. We find that Src42A activity is normally confined to the periphery of the border cell cluster. Loss-of-function, gain-of-function, and over-expression all affect the ability of the cells to complete their migration. Surprisingly, Src is also essential for specification of the proper number of migratory border cells. This establishes a genetically tractable, *in vivo* model for the study of the contribution of Src to collective cell migration in a complex environment.

## Results

### Src42A, not Src64B, is required for border cell migration

Of the two Src homologues in Drosophila, Src42A is more related to vertebrate Src, but the two proteins function redundantly in several cell types (Takahashi et al., 2005; Wouda et al., 2008). In the ovary, Src42A is abundantly expressed in all follicle cells [(Takahashi et al., 2005 and Figures 1A-B”), including border cells (Figures 1A-D”) and in germline cells, and the staining overlapped extensively with E-cadherin (Figure 1). In contrast, Src64B expression is restricted to the nurse cells (Figures 1E-G”)(O’Reilly et al., 2006).

**Figure 1.**
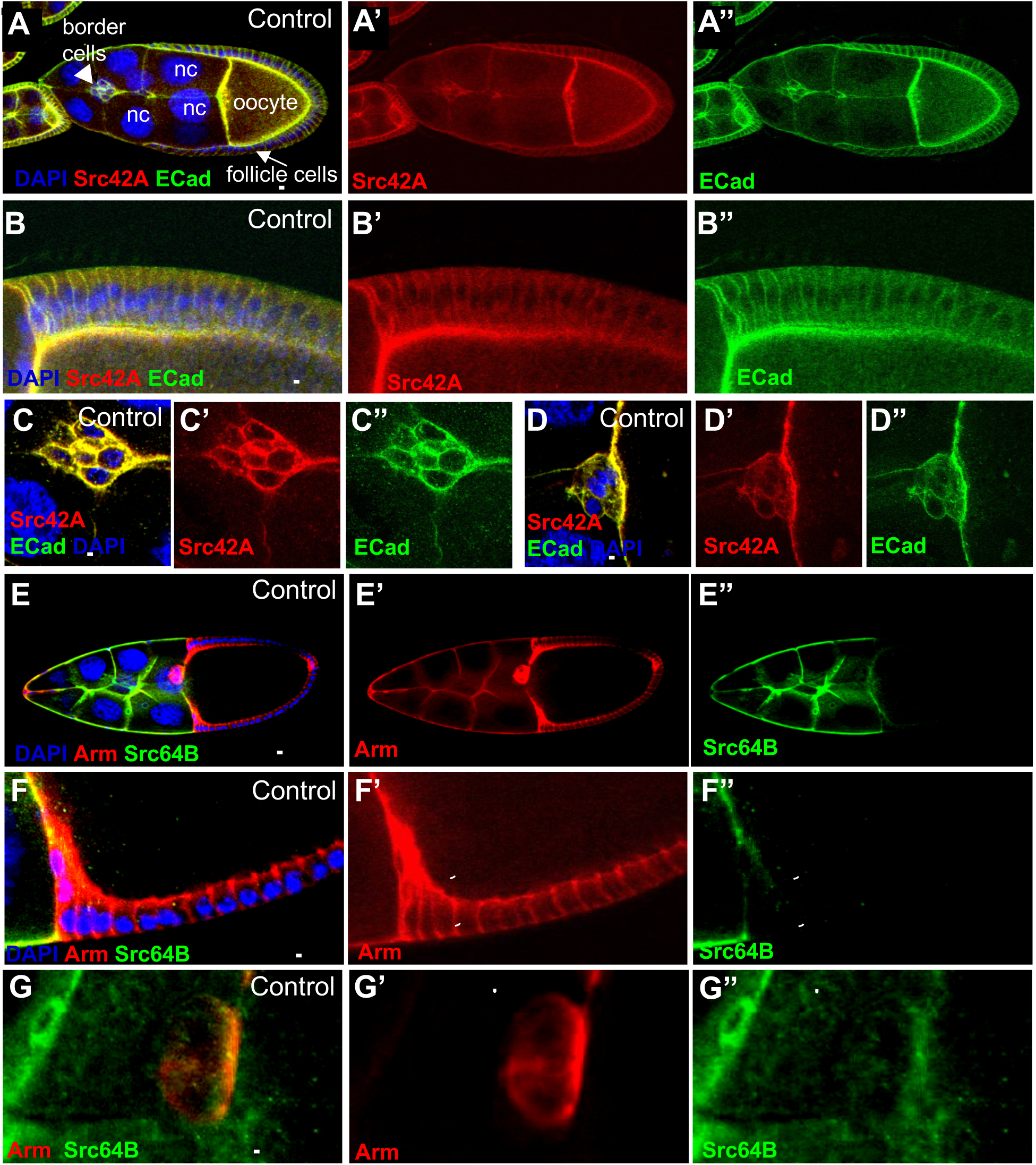
Src42A and Src64B expression patterns in egg chambers. Confocal images of (A-D”) Control (c306-Gal4; +; +) egg chambers stained with DAPI (blue), anti-Src42A (red), and anti-E-Cadherin (green). (A-A”) Stage 9. nc = nurse cells. (B-B”) High magnification view of follicle cells of a stage 10 egg chamber. (C-C" and D-D”) High magnification view of border cell clusters of egg chambers in stages 9 (C-C”) and 10 (D-D”). (E-G”) Control (c306-Gal4; +; +) egg chambers stained with DAPI (blue), anti-Armadillo (red), and anti-Src64B (green). (E-E”) Image of stage 10 egg chamber. (F-F”) High magnification view of follicle cells of a stage 10 egg chamber. The dashed line in F” indicates where Src64B is not detectable in the follicle cells stained with Armadillo shown in F’. (G-G”) High magnification view of a border cell cluster of a stage 10 egg chamber. The dashed line in G” indicates where Src64B is slightly detectable in the border cells stained with Armadillo shown in G’. Scale bars: 50μm (A-A”, E-E”) and 10μm (B-D”, F-G”).

Src42A null mutations are homozygous lethal, and the gene is located proximal to available FRTs making it impossible to investigate its function in mosaic clones. Therefore, we reduced Src42A expression using RNAi. Antibody staining showed that Src42A protein expression was undetectable in border cells from *c306-Gal4;UAS-Src42A RNAi* females (Figure 2A-B”). This Gal4 line was chosen for the RNAi experiments because of its early and strong expression in anterior (as well as posterior) follicle cells including all cells of the border cell cluster [both polar cells and outer migratory cells (Murphy and Montell, 1996)].

**Figure 2.**
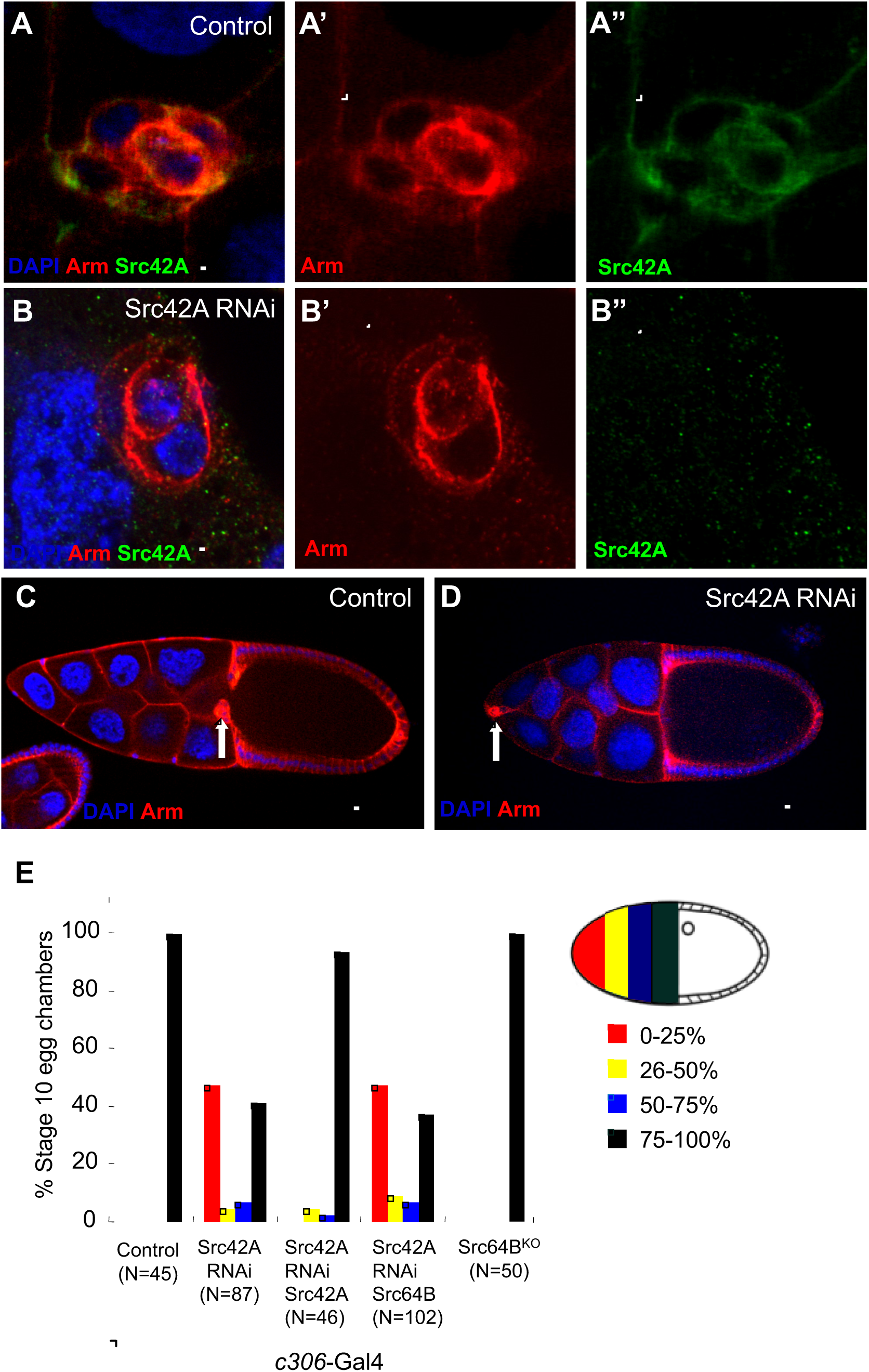
Src42A and not Src64B is required for border cell migration. (A-B) High magnification views of border cell clusters stained with DAPI (blue), anti-Armadillo (red), and anti-Src42A (green) in stage 9 egg chambers. (A-A”) A control border cell cluster (c306-Gal4;+;+). The dashed line in A” indicates where Src42A is detectable in the border cell cluster stained with Armadillo shown in A’. (B-B”) A border cell cluster of an egg chamber expressing UAS-Src42A RNAi (*c306*-Gal4; UAS-Src42A RNAi). The dashed line in B” indicates where Src42A is reduced in Armstained border cells shown in B’. (C) Control (*c306*-Gal4; +; +) and (D) Src42A knockdown (*c306*-Gal4; UAS-Src42RNAi) stage 10 egg chambers stained with DAPI (blue) and anti-Armadillo (red). Arrows in (C) indicates that the border cell cluster has completed migration and reached the oocyte. Arrow in (D) shows that the border cell cluster failed to detach from the anterior pole of the egg chamber. Scale bars: 10μm (A-B”) and 50μm (C-D). (E) Border cell migration defects caused by Src42A knockdown. Genotypes used were: 1) Control: *c306*-Gal4;+;+; 2) Src42A RNAi: *c306*-Gal4; UAS-Src42A RNAi; 3) Src42A RNAi/Src42A: *c306*-Gal4; UAS-Src42RNAi/UAS-Src42A; 4) Src42A RNAi/Src64B: *c306*-Gal4; UAS-Src42A RNAi/UAS-Src64B; 5) Src64B^KO^ is a homozygous viable Src64 null allele (O’Reilly et al., 2006).

The reduction of Src42A expression in border cells inhibited their migration such that 59% of clusters failed to reach the oocyte border by stage 10 (Figure 2C – E). This phenotype was rescued by co-expression of Src42A but not Src64B (Figure 2E), indicating that Src64B cannot substitute for Src42A during border cell migration. Src64B null mutants are viable (Dodson et al., 1998) and exhibit normal border cell migration (Figure 2E). Thus Src42A is required for border cell migration but Src64 is not.

### Src42A expression and activity levels are important in the outer, migratory cells

To assess the effects of manipulating Src activity levels, we expressed a kinase-dead version of Src42A, which may have dominant-negative effects (Src42A^DN^) and a constitutively active form Src42^CA^. These proteins caused lethality in combination with *c306*-Gal4, so we combined the temperature-sensitive repressor Gal80^ts^ together with *c306*-Gal4 and UAS-Src42A^DN^ and UAS-Src42^CA^. We raised the flies at 18°C to silence Gal4-dependent gene expression, and then shifted to 29°C (the non-permissive temperature for the repressor) four days prior to dissection. As expected, Src^DN^ caused a significant migration defect similar to the RNAi (Figure 3A, B, D). Over-expression of constitutively active (Src42A^CA^) or wild type (Src42A^WT^) kinases also caused significant migration delays (Figure 3C, D), suggesting that the levels of both expression and activity are important. Surprisingly, over expression of the wild type protein caused the most severe defect.

**Figure 3.**
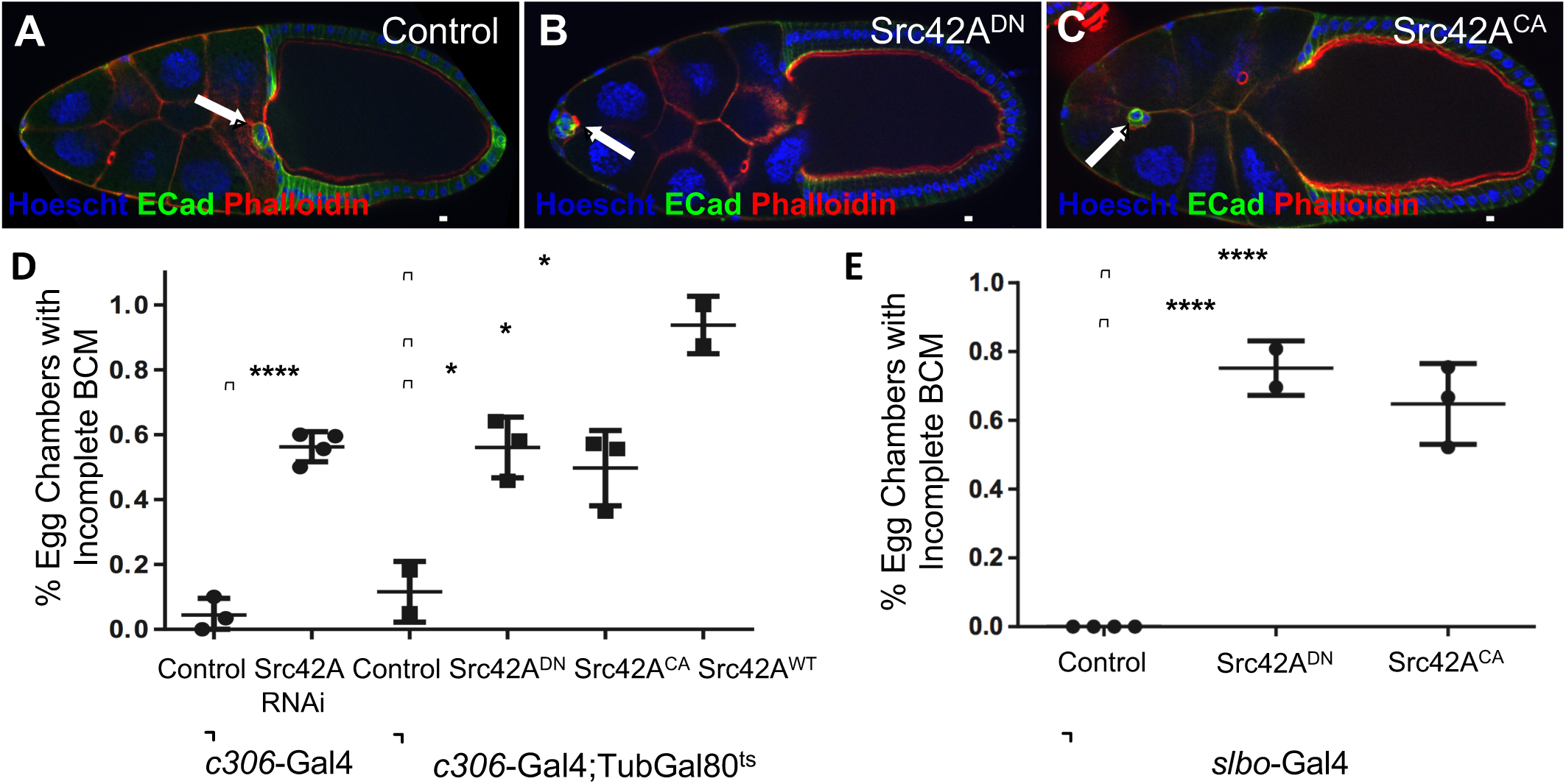
Incomplete border cell migration following Src42A loss‐ or gain-of-function. (A-C) Stage 10 egg chambers stained with Hoechst (blue), anti-E-cadherin (green), and Phalloidin (red). (A) Control (*c306*-Gal4/ w1118; +; +) border cell cluster completed migration. (B) Expression of kinase-dead Src42A (c306-Gal4; tubGal80^ts^/UAS-Src42A^DN^) in border cells caused detachment failure. (C) Src42A^CA^ (*c306*-Gal4; tubGal80^ts^/UAS-Src42A^CA^) expression impeded border cell migration. (D) Border cell migration defects in stage 10 egg chambers with(D) *c306*-Gal4 or (E) slbo-Gal4-driven Src constructs. Controls for (D) are *c306*-Gal4/w1118; +; + and *c306*-Gal4/w1118; tubGal80^ts^; +, respectively. Control for (E) is +; slbo-Gal4/cyo; +. P-values in (D) from left to right: <0.0001, <0.02, <0.04, and <0.02. P-values in (E) for both Src42A^DN^ and Src42A^CA^ <0.0001. Point representing Src42A RNAi in (E) represents an average of 78% incomplete border cell migration for three experiments. For (D) N = 15-60 and for (E) N = 15-45. Scale bar: μm (A-C).

C306-Gal4 is expressed in both non-migratory polar cells and outer, migratory cells of the border cell cluster. To determine if Src42A activity levels are important in the migratory cells, we used slbo-Gal4, which drives expression in outer, migratory cells but not polar cells (Geisbrecht and Montell, 2004) to drive Src42A^DN^ or Src^CA^. In both cases, migration was severely impaired (Figure 3E). Therefore, Src42A activity levels appear to be important autonomously in the migratory cells for their collective movement. Inhibition of Src42A expression in both polar and border cells with c306-Gal4 did not affect production or secretion of Upd from polar cells (Figure S1), further supporting the autonomous requirement for Src42A in outer, migratory cells.

### Src42A is required for border cell detachment

As border cells leave the anterior end of the egg chamber, they must detach from both the basal lamina that surrounds the egg chamber and the anterior follicle cells that stay behind. Src is known to destabilize cell-cell and cell-matrix adhesions (Wadhawan et al., 2011)(Frame et al., 2002). Therefore, we assessed the effect of Src42A inhibition on this detachment step. Inhibition of Src42A expression or activity caused a significant impairment in their ability to detach (Figure 4A-C). Intriguingly, the effect of RNAi was stronger than that of expressing the kinase-dead protein, suggesting that the kinase-dead protein may not function purely as a dominant-negative protein. The kinase-dead form of Src has previously been suggested to exhibit both dominant-negative and constitutively active characteristics (Read et al., 2004), which could be partially offsetting. RNAi is rarely completely effective in eliminating protein expression, so combining RNAi with heterozygous loss of function tends to enhance on-target (but not off-target) RNAi phenotypes. Combining the UAS-Src42ARNAi line with either of two different loss-of-function Src42A alleles in heterozygous condition, enhanced the border cell detachment defect (Figure 4C). We further confirmed the detachment defect by live imaging (Figure 4D-E).

**Figure 4.**
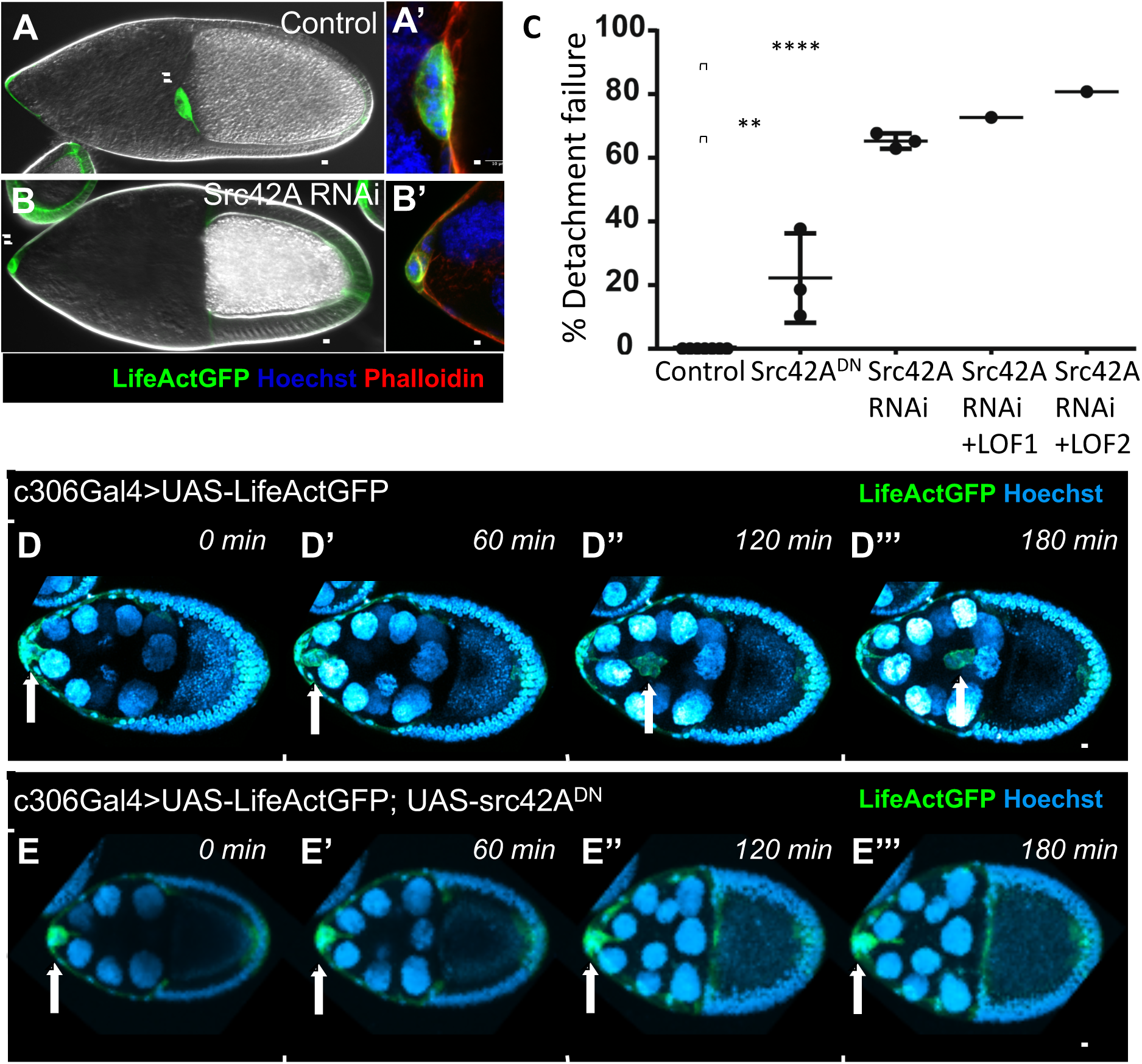
Failure of border cell cluster detachment following inhibition of Src42A. (A-B) Stage 10 egg chambers stained with Phalloidin (red) and Hoechst (blue), and express UAS-LifeActGFP under *c306*-Gal4; +; tubgal80^ts^ driver. (A, B) were visualized with DIC and (A’, B’) with fluorescent microscopy. Scale bar: 50μm (A-B) and 10μm (A’, B’). (A) Border cell cluster successfully detached from the anterior end of the egg chamber and completed migration by stage 10. (A’) High magnification image of border cell cluster at the oocyte border. (B) Border cell cluster fails to detach from the anterior end of the egg chamber and fails to migrate. (B’) High magnification image of border cell cluster at the anterior end of the egg chamber. (C) Plot showing percentage of border cell detachment failure in stage 10 egg chambers with *c306*-Gal4 driving Src42A under-expression phenotypes: UAS-Src42A^DN^, UAS-Src42A RNAi, UAS-Src42A RNAi with Src42A^E1^ loss-of-function allele, and UAS-Src42A RNAi with Src42A^myri^ loss-of-function allele. N=40-100. Statistical differences were found between the control and Src42A^DN^ (p-value<0.002) and Src42A RNAi (p-value<0.0001). (D-E) Time-lapse images from a 180 minute live-imaging experiment capturing border cell cluster detachment and border cell migration, stained with Hoechst (blue) and UAS-LifeActGFP expressed under *c306*-Gal4 driver. Arrows indicate border cell cluster. (D) Control egg chamber show border cell cluster successfully detaching from the anterior end of the egg chamber and migrating towards the oocyte border at the posterior end. (E) When Src42A is under-expressed in Src42A^DN^, the border cell cluster fails to detach from the anterior end of the oocyte and therefore cannot migrate. Scale bar: 50μm (D-E).

### Src42A is required for border cell specification

We noticed that early knockdown of Src42A with *c306*-Gal4 caused the border cell cluster to appear smaller than normal. Therefore, we counted the number of outer, migratory border cells and inner polar cells in *c306*-Gal4; UAS-Src42A-RNAi expressing clusters compared to control. We always observed two central cells with small nuclei, indicating normal polar cell development (Figure 5A-C). Whereas, slbo-Gal4 does not drive UAS-dependent gene expression in polar cells, the Slbo protein is normally expressed in polar cells and outer, migratory border cells and can be used as a marker [(Montell et al., 1992) and Figure 5]. In contrast to control clusters, which contain 4-7 (mean 4.5) outer, migratory cells, Src42ARNAi-expressing clusters contained between 0 and 6 (mean 2.5) (Figure 5D). Even those border cells that did form expressed lower levels of Slbo protein (Figure 5E).

**Figure 5.**
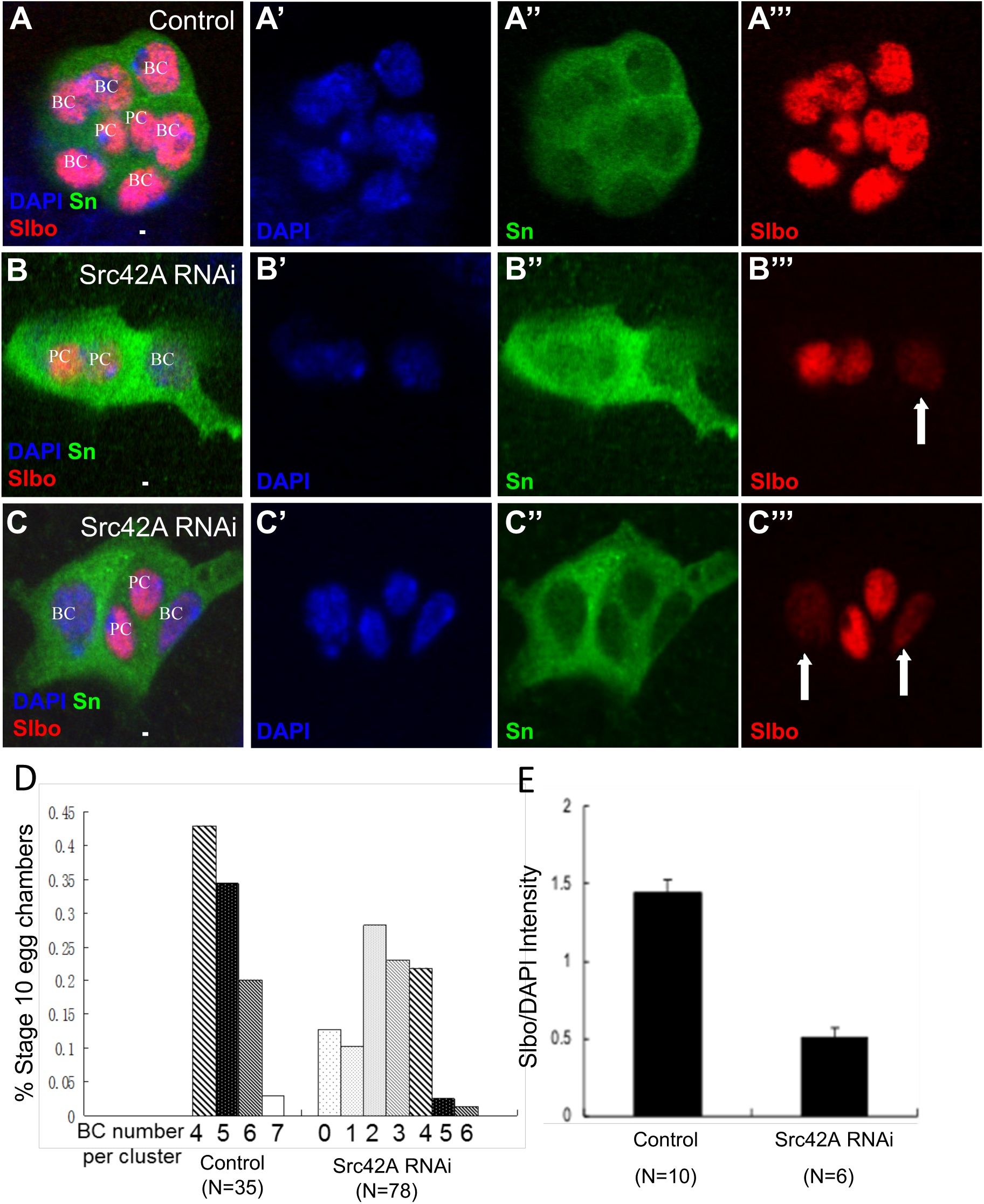
Src42A is required for the specification of migratory border cells. (A-C) High magnification view of border cell clusters stained with DAPI (blue), anti-Singed (SN, Green) and anti-SLBO (red). (A-A’”) A control (*c306*-Gal4;+;+) cluster with six outer border cells (BC) and two polar cells (PC) in the center, which express similar levels of Slbo. (B-C”) *Src42ARNAi* (*c306*-Gal4;UAS-Src42ARNAi;+) border cell clusters with two polar cells and only one (B-B”’, arrow in B”’) or two (C-C”’, arrow in C”’) outer border cell, which express less Slbo than the polar cells. (D) Quantification of the Slbo positive border cells number in control and UAS-Src42ARNAi egg chambers in stage 10. (E) Quantification of the ratio of Slbo nuclear staining to DAPI intensity for border cells in control and UAS-Src42ARNAi egg chambers.

### Src42A is active at the cluster periphery

The results described so far indicate that Src42A plays multiple roles in border cell development, including cell fate and differentiation as well as collective migration. Moreover, both the level of expression and activity appear to be critical. To gain insight into how Src activity might be regulated during border cell migration, we used an antibody that specifically recognizes the active form of Src (which is phosphorylated at tyrosine 419, equivalent to Y400 in Src42A) to stain egg chambers. In contrast to total Src, which co-localizes extensively with E-cadherin (Figure 1), active Src was enriched at border cell/nurse cell junctions, where E-cadherin levels are lowest. Border cell/border cell and border cell/polar cell junctions, where E-cadherin levels are higher lacked detectable active Src (Figure 6A, A’). Over-expression of Src42A^WT^, increased active Src both at the periphery and at border cell/border cell junctions (Figure 6B, B’, C).

**Figure 6.**
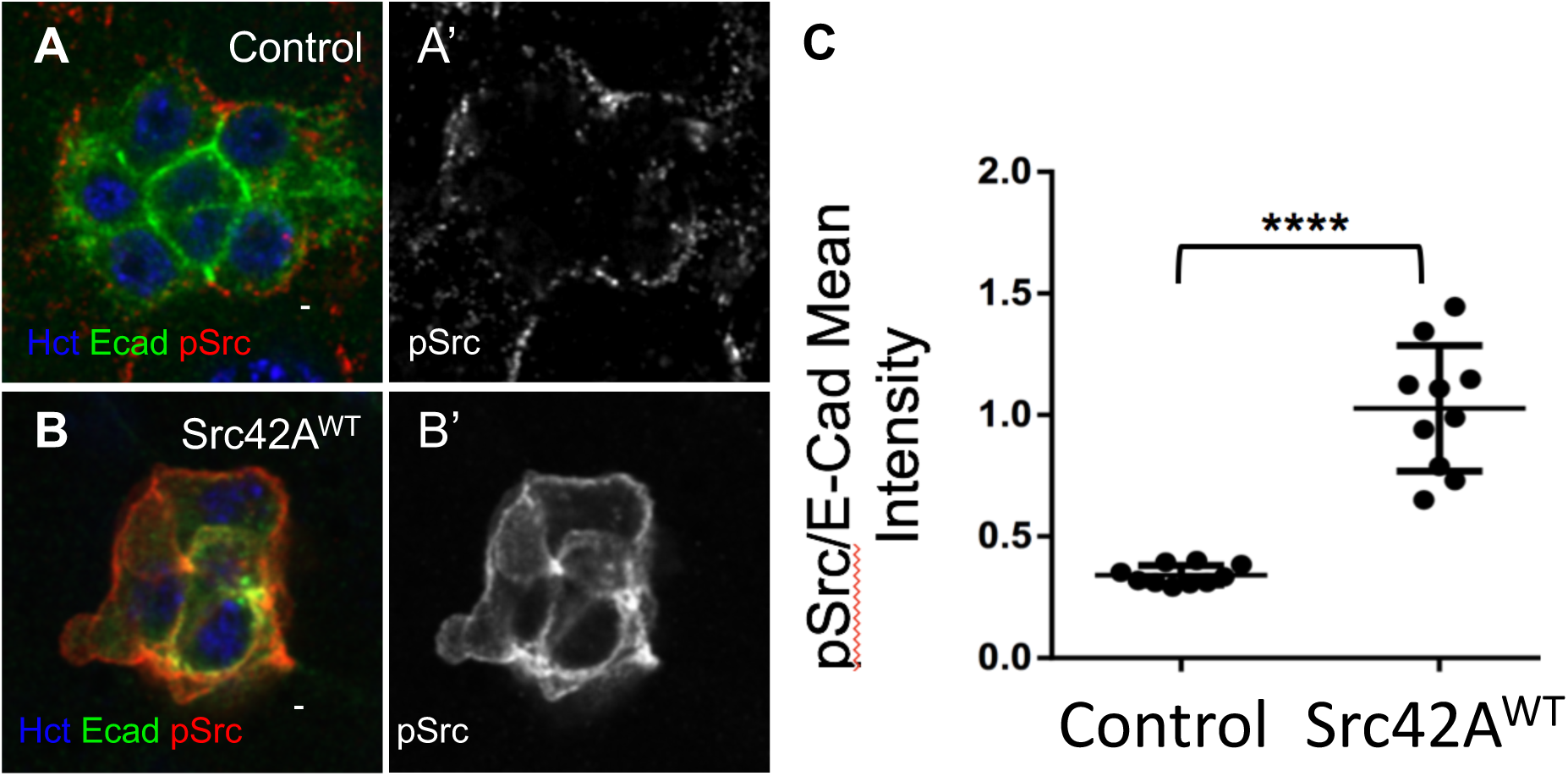
Activated Src42A levels in the border cell cluster are increased when Src42A is overexpressed. (A-B) Control (w1118; *slbo*-Gal4; +). and Src42A^WT^ (+; slbo-Gal4; UAS-Src42A^WT^) stage 9 egg chambers stained with anti-E-Cadherin (green), anti-pSrc (red) and Hoechst (blue). Scale bar: 10μm. (A’-B’) p-Src staining in A and B is shown in grey. (C) Quantification of anti-pSrc intensity normalized to anti-E-Cadherin expression in the border cell cluster. P-value <0.0001. N=10 egg chambers.

### Src42A^CA^ affects cell morphology and cluster cohesion

Border cell clusters are normally compact, which facilitates their collective movement (McDonald lab ref) (Figure 7A). Since Src kinases are known to inhibit cell-cell adhesion, we tested whether Src42^CA^ expression would inhibit adhesion within the cluster. Src42^CA^ expression was generally toxic. However, in those egg chambers that survived and developed to stage 9 or 10, expression of Src42^CA^ disrupted the compact organization of border cell clusters (Figure 7B-D). The cells filled with active Src and developed long, spindly protrusions (Figure 7E-F’). This abnormal morphology was also evident in centripetal cells (which express slbo-Gal4) as well (Figure 7G,G’) and has been noted previously (Somogyi and Rørth, 2004).

**Figure 7.**
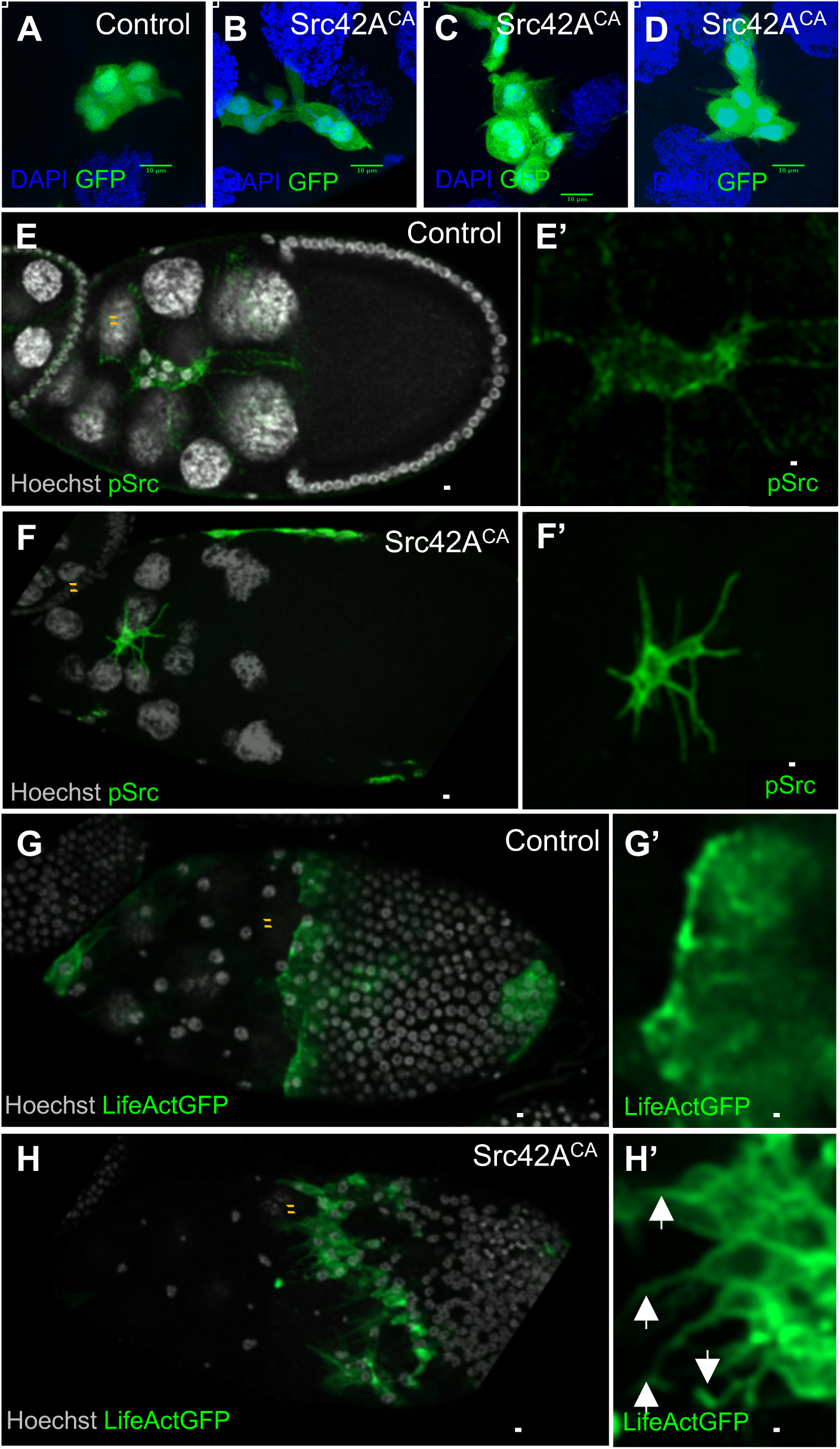
Loss of cluster cohesion and ectopic protrusions in Src42A^CA^-expressing cells. (A-D) High magnification images of border cell clusters in egg chamber stained with anti-GFP (green) and DAPI (blue). (A) Control border cells form a cohesive cluster. (B-D) *slbo*-Gal4 UAS-Src42A^CA^ border cells (E-I) Stage 10 egg chambers stained with anti-pSrc (green) and Hoechst (grey). (E) Control stage 10 egg chamber (w1118; *slbo*-Gal4; +) with (E’) high magnification of border cell cluster. (F) Ectopic protrusions in UAS-Src42A^CA^ expressing border cells (+; *slbo*-Gal4, *slbo*-LifeActGFP/UAS-Src42A^CA^; +). (F’) High magnification of a border cell with ectopic protrusions. (G) Control stage 10 egg chamber (w1118; slbo-Gal4, slbo-LifeActGFP; +) with (G’) high magnification of centripetal follicle cells. Centripetal follicle cells do not display ectopic protrusions. (H) Ectopic protrusions were observed in centripetal follicle cells when Src42A^CA^ was expressed (+; *slbo*-Gal4, *slbo*-LifeActGFP/UAS-Src42A^CA^; +). (H’) High magnification image of centripetal follicle cells displaying ectopic protrusions. Scale bar: 10μm (A-D, E’, F’, G’, H’) and 50μm (E-H).

## Discussion

Src is activated in numerous human cancers, and while much has been learned about the upstream signals and downstream targets of Src *in vitro*, little is known concerning its roles or mechanisms of action in collectively migrating cells *in vivo*. Here we show that Src42A is required for specification and collective migration of the border cells. Analysis of loss‐ and gain-of-function of Src allow us to conclude that Src42A but not Src64B is expressed, active, and required at several steps. First, Src42A is required to specify the correct number of border cells. Second, Src42A promotes the detachment of the cluster from the follicular epithelium and third, the limitation of the Src42A protein and activity is required in order to maintain the morphology and integrity of the cluster during collective cell migration. While previous studies have shown that constitutive activation of Src42A impedes border cell migration (Luo et al., 2015; Somogyi and Rørth, 2004), we show the requirement for, and the pattern of activity of, Src42A during this process.

Src proteins are composed of six conserved domains: an N-terminal myristoylated segment mediating attachment to cell membranes, an SH3 domain followed by an SH2 protein-protein interaction domain, a linker domain, a tyrosine kinase domain and a C-terminal regulatory segment. This functional domain architecture allows Src to play roles through its kinase activity and also by interacting with a variety of proteins and signaling pathways, raising the question as to how Src kinase activity is regulated and what the downstream effectors of Src are in collective border cell migration. Our results suggest that there may be novel mechanisms of action to be uncovered for Src in the specification and collective migration of the border cells.

### Regulation of Src expression levels and activity are essential

Of the many upstream signals that can in principle activate Src tyrosine kinases, it seems most likely that receptor tyrosine kinases (RTKs) activate Src42A in border cells, though this remains to be established. PDGF‐ and VEGF-related receptor (PVR) and fly EGF receptor (EGFR) are required for border cell migration. The ligands are produced by the germline (nurse cells and/or oocyte)(Duchek and Rørth; McDonald et al.,(Duchek and Rørth, 2001; Duchek et al., 2001; McDonald et al., 2003, 2006), therefore RTK activity in border cells is likely to be highest at nurse cell/border cell interfaces where we have found that Src activity is highest. Our results show that active Src is likely rapidly dephosphorylated or degraded in border cells, as in other cell types (Harris et al., 1999), because the level of active Src is normally challenging to detect. Over-expression of Src^WT^ results in a large increase and ectopic localization of active Src in border cells, suggesting that negative regulators are limiting. One such negative regulator appears to be the receptor for activated C kinase, Rack (Luo et al., 2015), though its mechanism of activation and action remain to be determined.

### The role of Src in border cell migration is likely unrelated to focal adhesion turnover

We found that Src42A is essential for border cell specification, detachment, and migration. Border cells detach from a basal lamina rich in type IV collagen and laminin as they initiate migration. They then migrate in between nurse cells. While there is strong evidence that dynamic E-cadherin-mediated adhesion between border cells and nurse cells is required for border cell migration (Cai et al., 2014; Niewiadomska et al., 1999), there is no evidence for cell-matrix adhesion playing a significant role in this setting. While there is abundant evidence for Src and focal adhesion kinase (FAK) regulating each other and promoting focal adhesion turnover in fibroblasts migrating on fibronectin-coated coverslips, this mechanism is unlikely to contribute to border cell migration. Drosophila FAK is not required for border cell migration (Grabbe et al., 2004). Src-dependent focal adhesion turnover may contribute to border cell detachment from the basal lamina, however if so, the mechanism does not appear to depend on FAK. Another downstream target of Src implicated in cell migration is cortactin; however cortactin causes a mild defect in border cell migration (Somogyi and Rørth, 2004). While focal adhesion turnover is thus not the most likely effect of Src on border cell migration, adherens junction turnover is a more promising candidate. Fly Src colocalizes with adherens junction proteins in numerous tissues, including follicle cells generally and border cells in particular (Takahashi et al., 2005 and Figure 1). Yet active Src localizes most strongly in nurse cell/border cell interfaces (this study), where E-cadherin is active and required but does not accumulate to high levels. Thus, one model is that Src is activated by RTK signaling specifically at border cell/nurse cell boundaries, resulting in internalization of E-cadherin allowing for the dynamic border cell/nurse cell adhesion required for migration.

### The role of Src in border cell specification may be independent of STAT

The only signal known to be required for border cell fate specification to date is Upd/Jak/STAT. Since polar cells are the source of Upd, the highest Jak/STAT activity at stage 9 is normally restricted to border cells (and posterior follicle cells). Multiple feedback mechanisms establish a threshold so that cells with the highest Jak/STAT activity acquire border cell identity, express SLBO, and migrate while cells that experience lower and/or transient Jak/STAT assume so-called stretch cell identity and remain within the epithelium (Starz-Gaiano et al., 2008; Yoon et al., 2011). Ectopic expression of Upd or activated Hop is sufficient to specify ectopic migratory border cells (Silver and Montell, 2001; Silver et al., 2005). Since Src kinases are known to cooperate with Jak to activate mammalian STATs, and blocking STAT activity strongly suppresses Src-dependent overgrowth in a Drosophila tumor model (Garcia et al., 2001; Read et al., 2004; Ren and Schaefer, 2002; Zhang et al., 2000), it seems logical to propose that the border cell specification defect that we observe upon inhibition of Src42A might be due to defective STAT activation. Yet we have been unable to detect a reproducible effect of Src inhibition on STAT accumulation. If this is the case, it suggests that an additional Src-dependent pathway, independent of STAT, is required for border cell specification.

Due to the genetic tractability of the Drosophila border cell system and its amenability to live imaging, this work establishes an important model for unraveling the regulation and function of Src in collective cell behavior *in vivo*. Given the emerging importance of collective cell migration to tumor metastasis (Aceto et al., 2014; Cheung and Ewald, 2016; Cheung et al., 2016; Lambert et al., 2017), insights derived from border cells may be critical in unraveling the still-mysterious contributions of Src to tumor progression. While the importance of collective cell migration for tumor metastasis is increasingly appreciated, cell-on-cell migration is still studied far less than cells migrating on or through extracellular matrix. So far, border cell studies suggest that the mechanisms by which Src regulates collective cell-on-cell migration may well differ from the most commonly studied functions of Src in individual, cell-on-matrix migration.

### Experimental Procedures

#### Fly stocks and genetics

UAS-Src42A RNAi line is from the Vienna Drosophila Resource Center (construct ID: 108017). UAS-Src42A^DN^ and UAS-Src42A^CA^ are from Dr. Tetsuya Kojima. UAS-Src42A and UAS-Src64B are from Dr. Tian Xu. Src64B null allele is from Dr. Alana O’Reilly. Src42A LOF1 (Src42A^E1^ allele, stock number: 6408) and LOF2 (Src42A^myri^ allele, stock number: 6453) mutant alleles are from the Bloomington Drosophila Stock Center.

The drivers used in this study were *c306*-Gal4 (Murphy and Montell, 1996) and slbo-Gal4 (Rorth et al., 1998, Geisbrecht and Montell, 2004). UAS-GFP was used for Figure 6D-G. Crosses were carried out at 25°C unless noted otherwise. For Figures 1, 2, 3E, 5, 6, and 7A-D, 1-day old flies were fattened at 29°C for 20-24 hours before dissection. For Figures 3A-D, 4, and 7E-H, 1-day old progeny were moved to 29°C for 3 days total and were fattened for 20-24 hours before dissection. For the experiments including tubGal80^ts^, the crosses were carried out at 18°C and the progeny moved to 29°C for 4 days total and were fattened for 20-24 hours before dissection. Typical controls are the driver lines crossed to w^1118^.

### Immunofluorescence

Ovaries were dissected in Schneider’s medium (Thermo Fisher Scientific, Waltham, MA) with 20% fetal bovine serum. For Figures 1, 2, 3E, 5, and 7A-D, ovaries were fixed in 4% formaldehyde (Polysciences, Inc.) for 20 minutes, washed three times in PBT buffer (1X PBS, 0.2% Triton), and then blocked in PBT plus 5% normal goat serum (Sigma-Aldrich) for 30 min. For Figures 3A-D, 4, 6, and 7E-H, ovaries were fixed in 4% paraformaldehyde (Electron Microscopy Sciences), and washed in PBT (1X PBS, 0.3% Triton) without goat serum. Egg chambers were incubated in PBT buffer containing primary antibodies for 2 hours at room temperature or overnight at 4°C, washed 3 times in PBT for 20 minutes each, then incubated in PBT containing fluorescence-conjugated secondary antibodies for 2 hours at room temperature. The following antibodies were used: mouse anti-Armadillo N27A1 (1:100, DSHB), anti-E-cadherin (1:50, DSHB DCAD2), mouse anti-Singed (1:25, DSHB), rabbit anti-GFP (1:500, Invitrogen), rat anti-SLBO (1:5000, from Pernille Rørth), rabbit anti-Src42A (1:500, gift from Tetsuya Kojima), rabbit anti-pSrc 2101S Tyr416 (1:25, DSHB), and anti-Upd. The secondary antibodies used were Alexa Fluor^®^ 488, 568, and 647 (1:200 or 1:400, Invitrogen). Alexa Fluor^®^ 488 and 568 Phalloidin were used to detect filamentous actin (1:400 or 1:200, Thermo Fisher). DNA was visualized with Hoechst (1:1000, Invitrogen) or DAPI (1:1000, Sigma-Aldrich). All tissues were mounted in Vectashield (Vector Laboratories, Burlingame, CA).

### Egg chamber culture and live imaging

Ovarioles were dissected and cultivated according to our published protocol, including insulin at 40μg/ml^-1^. Border cell migration was followed in real time by the expression of UAS-LifeActGFP under the control of *c306*-Gal4 driver for Figure 4E-F, and *slbo*-Gal4 driver for Figure 6. For Figure 4E-F, Hoechst was added to the culture medium for DNA visualization.

### Image acquisition and treatment

Images for Figures 1, 2, 5, and 7A-D were taken with Zeiss 510 MicroImaging Confocal Laser Scanning Microscope using a 25x or 63x objective lens, along with LSM 510 AIM acquisition software. Images for Figures 3 and 4A-B were taken with Zeiss Axio Imager. M2 with ApoTome using a 40X objective lens. Images for Figures 4D-E, 6, and 7E-H were taken with Zeiss LSM800 Confocal microscope, using 20X and 40X objectives, using Zen acquisition software. Image Z-stacks were exported in their original format and processed using Fiji software.

### Statistical analysis

We used GraphPad Prism 6 software for unpaired two-tailed t-test statistical analysis in Figures 3, 4, and 6.

## Acknowledgements

We thank Jingchuan Luo for confirming some of the results presented in this manuscript. This work was supported by NIGMS grant GM73164 to DJM.

---

**Supplementary Figure 1.**
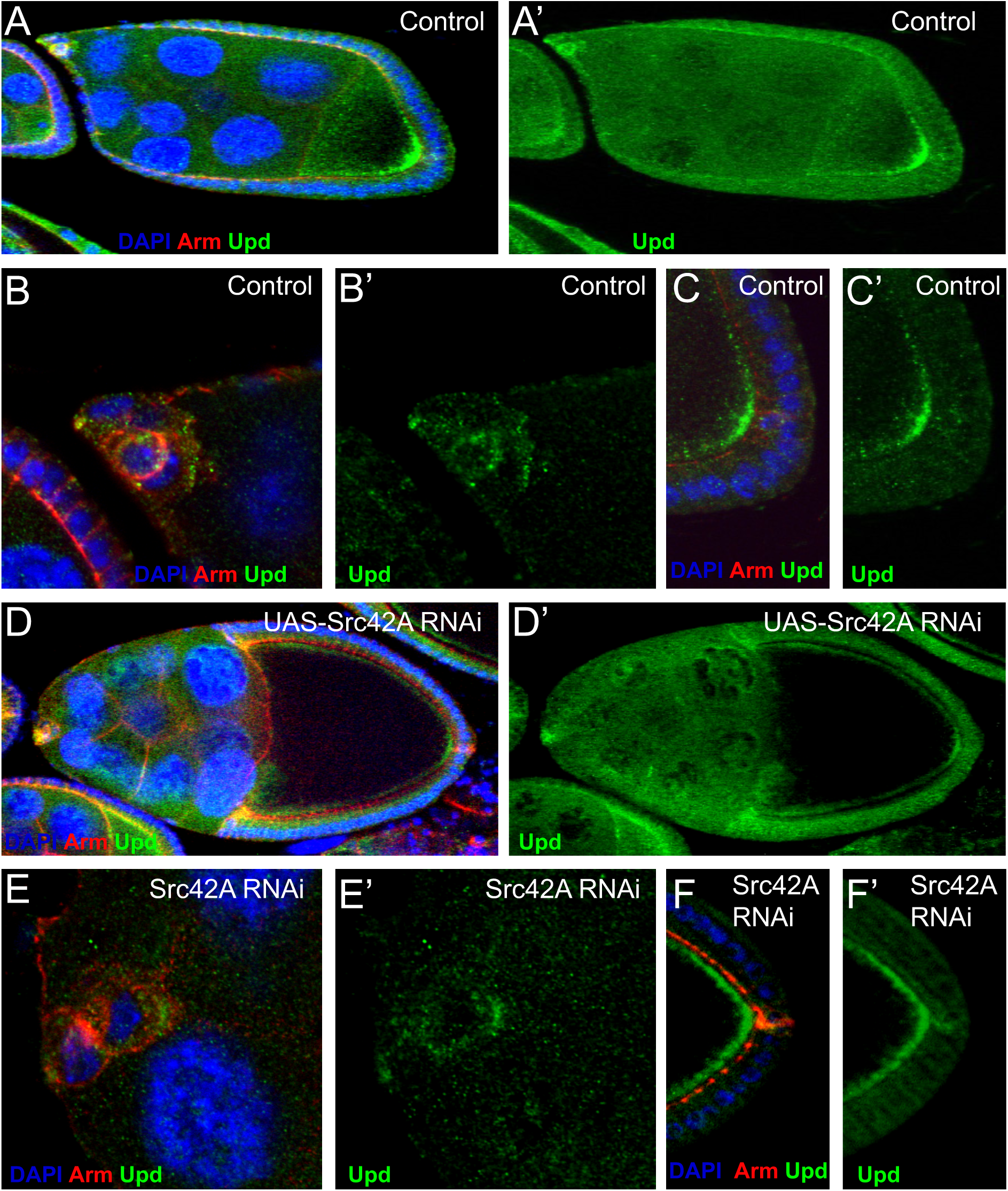
Upd expression was unaffected by Src42ARNAi. (A) A wild type stage 9 egg chamber stained with DAPI (blue), anti-Arm(red) and anti-Upd (green). Anterior is to the left. Upd is enriched around anterior polar cells (arrow) and in a gradient emanating from the posterior polar cells (open arrow). (A’) Anti-Upd staining only. (B, C) High magnification views of wild-type border cells (B) and posterior follicle cells (C) stained with DAPI (blue), anti-Arm (red) and anti-Upd (green). (B’ and C’) Corresponding anti-Upd staining only. (D-D’)A late stage 9 egg chamber from *c306*-Gal4;UAS-Src42ARNAi stained with DAPI (blue), anti-Arm(red) and anti-Upd (green). E-F’) High magnification views of a *c306*-Gal4; UAS-Src42ARNAi egg chamber stained with DAPI (blue), anti-Arm (red) and anti-Upd (green). (E, E’) Border cells. (F, F’) Posterior follicle cells.

